# Rationally designed Gla-domainless FXa as TFPI bait in hemophilia

**DOI:** 10.1101/2022.08.03.502629

**Authors:** Marie-Claire Dagher, Atanur Ersayin, Landry Seyve, Mathieu Castellan, Cyril Moreau, Luc Choisnard, Nicole Thielens, Raphaël Marlu, Benoît Polack, Aline Thomas

## Abstract

Gla-domainless factor Xa (GD-FXa) was proposed as a trap to the endogenous anticoagulant Tissue Factor Pathway Inhibitor (TFPI) to restore thrombin generation in hemophilia. Using computational chemistry and experimental approaches, we previously showed that S195A GD-FXa also binds TFPI and restores *ex vivo* coagulation in hemophilia plasmas.

To design a GD-FXa variant with improved anti-TFPI activity and identify suitable sites for mutagenesis, we performed molecular dynamics simulations. The calculations identified residues R150_FXa_ and K96_FXa_ as cold-spots of interaction between GD-FXa and the K2 domain of TFPI. In the three-dimensional model, both residues are facing TFPI hydrophobic residues and are thus potential candidates for mutagenesis into hydrophobic residues to favor an improved protein-protein interaction.

Catalytically inactive GD-FXa variants containing the S195A mutation and additional mutations as K96Y, R150I, R150G and R150F were produced to experimentally confirm these computational hypotheses. Among these mutants, the R150F_FXA_ showed increased affinity for TFPI as theoretically predicted, and was also more effective than S195A GD-FXa in restoring coagulation in FVIII deficient plasmas. Moreover, the R150 mutants lost interaction with antithrombin, which is favorable to extend their half-life.

## Introduction

Hemophilia is a bleeding disorder due to the deficiency of either factor VIII or IX (FVIII or FIX) which decreases the production of thrombin from the intrinsic pathway of coagulation. Substitution of FVIII or FIX by plasmatic or recombinant proteins is therapeutically efficient but can lead to the generation of neutralizing antibodies. Recombinant FVIIa (Novoseven ®) or aPCC (FEIBA®) are approved bypassing agents that can be administered to restore thrombin generation in patients with inhibitors^1^. Clinical trials on hemophilia have recently been reviewed^2^. The development of extended half-life replacement factors is one strategy to decrease the frequency of factor injections and to improve the quality of life of patients^3^. Non-factor therapies have been developed such as Emicizumab, (ACE910; Roche), a bispecific antibody mimicking FVIIIa and simultaneously binding FIXa and FX^4^; it is approved in USA and Europe for the routine prophylaxis of bleeding episodes in hemophilia A. Among the non-factor replacement therapies under development^3, 5^ some aim at targeting endogenous anticoagulants such as antithrombin, activated protein C and TFPI^6^. Fitusiran, a siRNA against antithrombin, is currently in clinical phase III^7^.

Physiologically, the complex combining activated factor IX (FIXa) and its cofactor, factor VIIIa (FVIIIa) generates factor Xa (FXa) which, in cooperation with factor Va, activates prothrombin to thrombin leading to subsequent conversion of soluble fibrinogen into fibrin fibers. In hemophiliacs, small amounts of FXa can still be produced by the extrinsic tenase complex made of factor VIIa (FVIIa) and Tissue Factor. However, these two proteins are inhibited by TFPI. There are two major isoforms of TFPI: TFPIα, which contains three Kunitz-type domains (K1, K2, K3) followed by a positively charged C-terminus tail, and TFPIβ that contains the same K1 and K2 domains, followed by a C-terminus tail with a glycosylphosphatidylinositol anchor. Structurally, TFPI inhibits respectively FVIIa and FXa by inserting its Kunitz domain K1 and K2 in each active site cavities of these coagulation factor^8^. Relieving the TFPI inhibition of FXa is a means to restore coagulation, and some teams have developed monoclonal antibodies^9–12^ (among them Concizumab®^13^, ^14^, that benefits of a compassionate use program), aptamers^15^, and peptides^16^ as baits to TFPI. We have developed Gla-domainless factor Xa (GD-FXa), a FXa variant deprived of Gla domain^13^. This engineered GD-FXa shows procoagulant properties but some instability. When the catalytic serine S195 is mutated into an alanine, the derived S195A GD-FXa is still able to bind to TFPI, as predicted by our molecular dynamics studies^17^; this catalytically inactive GD-FXa shows improved stability and restores thrombin generation in hemophilia plasmas^17^. Portola Pharmaceuticals have commercialized it (andexanet alpha, Ondexxya®) in Europe and the United State, where it is used as an antidote to the Direct Oral Anticoagulants Rivaroxaban and Apixaban that target FXa.

We followed a computational chemistry approach to rationally design S195A GD-FXa variants with improved anti-TFPI activity. The complexes between GD-FXa (both wild-type and catalytically inactive) and the K2 domain of TFPI were modeled and submitted to a 200 ns molecular dynamics (MD) simulation. This allowed the identification of GD-FXa residues showing positive interaction energies with TFPI (i.e. GD-FXa residues contributing to the destabilization of the interaction with the K2 domain). These residues identified as cold-spots of interaction are good candidates for mutation to engineer an optimized bait to TFPI.

Previous MD simulations predicted that the catalytically inactive S195A GD-FXa mutant binds TFPI with an affinity similar to that of wild-type GD-FXa, which was experimentally validated^17^. Following a similar protocol, we present herein the identification of GD-FXa cold-spots of interaction with TFPI based on theoretical calculations, notably residues K96_FXa_ and R150_FXa_. A series of five GD-FXa mutants at positions K96 and R150 are produced in the S195A context. These mutants are functionally characterized, using thrombin generation assays and surface plasmon resonance (SPR) to respectively quantify their pro-coagulant activity and their ability to bind to two physiological inhibitors of the coagulation cascade: TFPI and antithrombin. We show that the R150F mutant has a significantly enhanced TFPI binding capacity and an improved pro-coagulant activity at low concentration; we show that this mutant escapes from antithrombin interaction which is a prerequisite for a future novel hemophilia treatment.

## Methods

### Molecular Dynamics

The computational procedure is fully described in the Supplementary Material. Briefly, we modeled the complex between the catalytic domain of FXa and the K2 domain of TFPI (TFPI-K2) and mutated the catalytic S195 into Alanine. The resulting model was subjected to 200 ns MD in explicit water, using CHARMM^18, 19^.

### Production of S195A GD-FXa mutants and TFPIα

The cDNA sequences (Genscript) were used for expression in Drosophila S2 cells (Life Technologies) using the plasmid pIEX (Merck Millipore) with a MB7 signal sequence. GD-FXa was produced as a single chain, the activation peptide replaced by a linker containing a double Furin cleavage site^20^. An HPC4 tag (Protein C derived epitope) was set at the C-terminus. S2 cells were transfected (Freestyle Max Reagent, Life Technologies); the supernatant was collected at Day 6 post-transfection. TFPI, purified on SP Sepharose column, displayed 95% purity. GD-FXa was very unstable and could not be purified. After cation exchange chromatography, S195A-GD-FXa protein was further purified on an anti-Protein C Affinity matrix (Roche). Maldi mass spectrometry showed that TFPI was glycosylated. Its functionality was assessed by inhibition of FXa activity on the chromogenic substrate PNAPEP1025 (Cryopep) and by binding to FXa by SPR.

The mutants S195A GD-FXa with an additional mutation namely R150G, R150I, R150F, K96Y, and the triple mutant K96YR150FS195A, were produced using the Quickchange procedure.

### Surface plasmon resonance spectroscopy

Analyses were performed at 25 °C using a Biacore 3000 instrument (GE Healthcare). The interaction of the mutants with immobilized TFPI (1500 to 2100 RU) was analyzed as described previously^17^. The kinetic constants and the resulting apparent equilibrium dissociation constants (*K*_D_ = *k*_d_/*k*_a_) were obtained by fitting the binding curves to a Langmuir 1:1 model using the Biaevaluation 3.2 software. The concentrations of the mutants varied from 1.25 to 40 nM.

AT (Cryopep) (diluted to 6 μg/mL in 10 mM sodium acetate pH=5) was immobilized on a CM5 sensor chip using the amine coupling chemistry in HBS-P buffer. The immobilization level ranged from 2600 to 4400 RU. Binding of GD-FXa mutants was measured at a flow rate of 20 μl/min in 18 mM Hepes, 135 mM NaCl, 2.5 mM CaCl2, 0.005% surfactant P20, pH 7.35. The specific binding signal was obtained by subtracting the background signal over the reference surface. Steady state analysis was used to calculate the *K*_D_ of the mutants. The binding responses at equilibrium obtained at six mutants concentrations ranging from 31.2 nM to 1 μM were plotted against the GD-FXa concentrations and *K*_D_ were calculated.

### Thrombin generation assays

Thrombin generation was assayed in severe hemophilia A plasma (Cryopep)^17^. Plasma was spiked with S195A GD-FXa or with the mutant proteins. Recombinant FVIII (Octocog alpha, Bayer Pharma) was used as positive control. Thrombin generation was triggered by 1pM tissue factor and 4μM phospholipids (PPP-reagent low, Diagnostica Stago) in the presence of a fluorogenic substrate (FluCa kit). All assays were run in triplicates on a CAT fluorimeter (Diagnostica Stago). Thrombin generation was monitored (excitation at 390 nm, emission at 460 nm for 60 min, 37°C).

## Results

### Molecular Dynamics

The analysis of the 200 ns MD simulation of the complex between S195A GD-FXa and TFPI-K2 allowed to quantify the distribution of interaction energies between each GD-FXa residue and the whole TFPI-K2 chain along MD (Table S1). The interaction energy, corresponding to the sum of the van der Waals and electrostatic energies, is an indicator of the stability of a complex.

Some GD-FXa residues show very low interaction energies with the TFPI-K2 chain (see Table I) as they strongly contribute to the interface stabilization: notably residue D189_FXa_ shows an interaction energy equal to E = −60.61 ± 0.36 kcal/mol, reflecting the fact that it is strongly hydrogen-bonded to R15_K2_ along the dynamics. Similarly, residues E37_FXa_ and E39_FXa_ show very stabilizing interaction energies respectively equal to E = −64.47 ± 1.58 kcal/mol and E = −78.34 ± 2.18 kcal/mol. During the whole MD trajectory, E37_FXa_ and E39_FXa_ entertain stable hydrogen bonds with respectively R32_K2_ and K34_K2_.

Some GD-FXa residues show small positive interaction energies along MD, and thus destabilize the protein-protein interface; these cold-spots of interactions are good candidates to design mutants with improved affinity for TFPI-K2. The distribution of residual interaction energies along the MD (Table S1) allows to identify three of these candidate residues, namely N35_FXa_, K96_FXa_ and R150_FXa_. The side-chains of these residues are located in spatial proximity to TFPI-K2 in the complex model, and are neither buried nor involved into local hydrogen bond network. However, N35G_FXa_ is located between two strong favorable interactions namely E24_FXa_-K34_K2_ and E22_FXa_-R32_K2_, therefore, it was not considered suitable for mutation in order to maintain those interactions.

On the contrary, residues K96_FXa_ and R150_FXa_ can be mutated without destabilizing the local internal hydrogen bond network. Residue K96GD-FXa gives rise to a small unfavorable energetic contribution and was considered as a potential mutation-site as it is facing L39_K2_; we replaced the long basic chain of K96 by a bulky hydrophobic tyrosine to favor local hydrophobic interactions. Residue R150_FXa_ was also selected as a good candidate for mutagenesis as it is located in a flexible loop and its side chain is pointing towards the solvent along the whole dynamics; furthermore, it is located less than 5 Å of the hydrophobic residue Y17_K2_; we hypothesized that replacing R150_FXa_ by a hydrophobic residue could be favorable for the interaction with the neighboring Y17_K2_.

It is to be noted that residue R150_FXa_ is not involved in protein-protein interfaces within the ternary complex model TF-FVIIa-FXa ^21^ nor within the prothrombinase model^22^. However, it is located at an exosite that is involved in the interaction with antithrombin (AT), as can be seen in the crystal structure of the ternary complex between AT, S195A FXa and the pentasaccharide fondaparinux (PDB ID: 2GD4^23^) (Figure 1). In this complex, residue R150_FXa_ entertains three side-chain hydrogen bonds with M251_AT_, E237_AT_ and N233_AT_. Thus, the replacement of R150_FXa_ by a hydrophobic residue, cancelling these favorable interactions, should very likely prevent the derived FXa mutants to be trapped by antithrombin, which could also favor the expected pro-coagulant activity.

**Figure 1:**
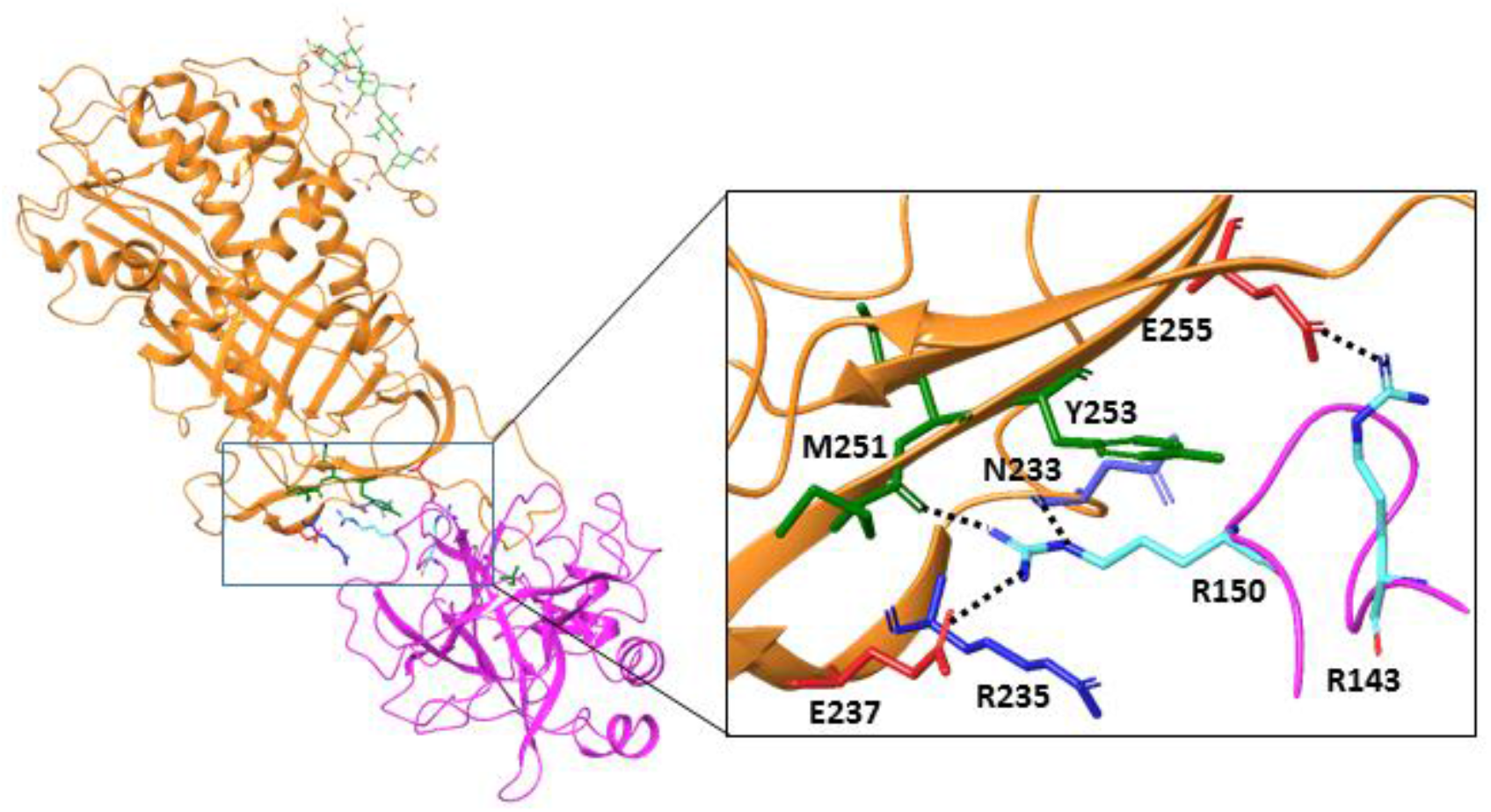
Structural analysis of the role of R150 in AT binding to FXa. Representation of the crystallographic structure of the complex between antithrombin and S195A FXa (respectively in orange and magenta ribbons), in the presence of pentasaccharide (in green sticks) (PDB ID: 2GD4^23^). The R150_FXa_ side chain is stabilized by three hydrogen bonds involving AT residues E237, E251 and N233.

### Biological validation of computer predictions

#### SPR measurement of affinity constants of GD-FXa mutants with TFPI

The different mutations at positions 96 and 150 were introduced in the catalytically inactive S195A GD-FXa. The mutants were produced in S2 cells and their concentration and purity were analyzed by SDS-PAGE and the Imagelab software (Bio-rad) see Figure S1. Their interaction kinetics with immobilized TFPI were analyzed as described^17^ (see representative binding data sets for each mutant in Figure S2). As predicted by MD calculations, R150FS195A GD-FXa shows a significantly lower association rate constant (*k*_a_) than S195A GD-FXa (see Figure 2), but since it also has a lower dissociation rate constant (*k*_d_), the resulting *K*_D_ is similar to that of S195 GD-FXa. The K96Y mutation was not favorable, resulting in an increased dissociation constant. When associated with the R150F mutation in the triple mutant, it counteracted the effect of the R150F mutation on the association constant.

**Figure 2:**
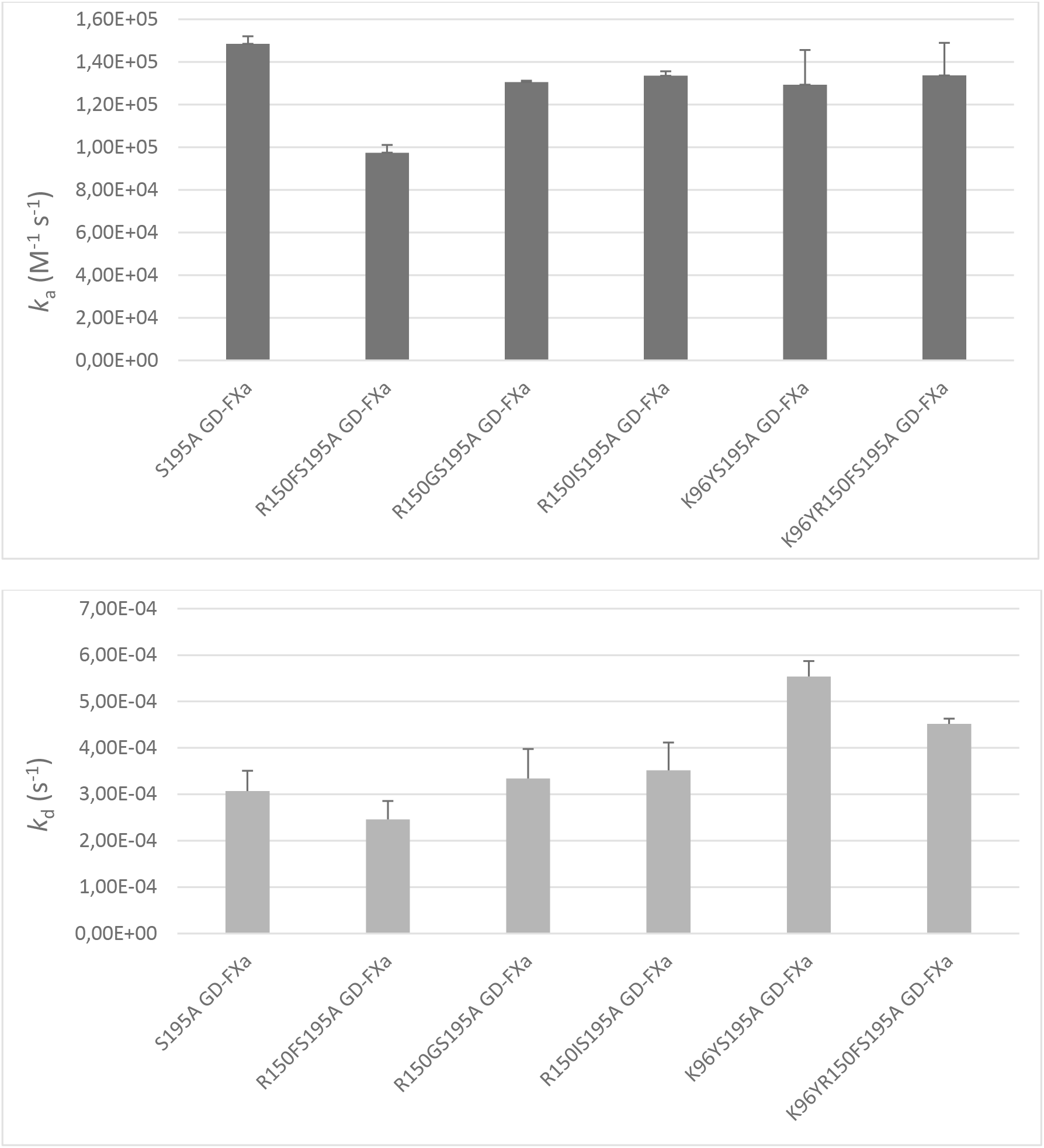
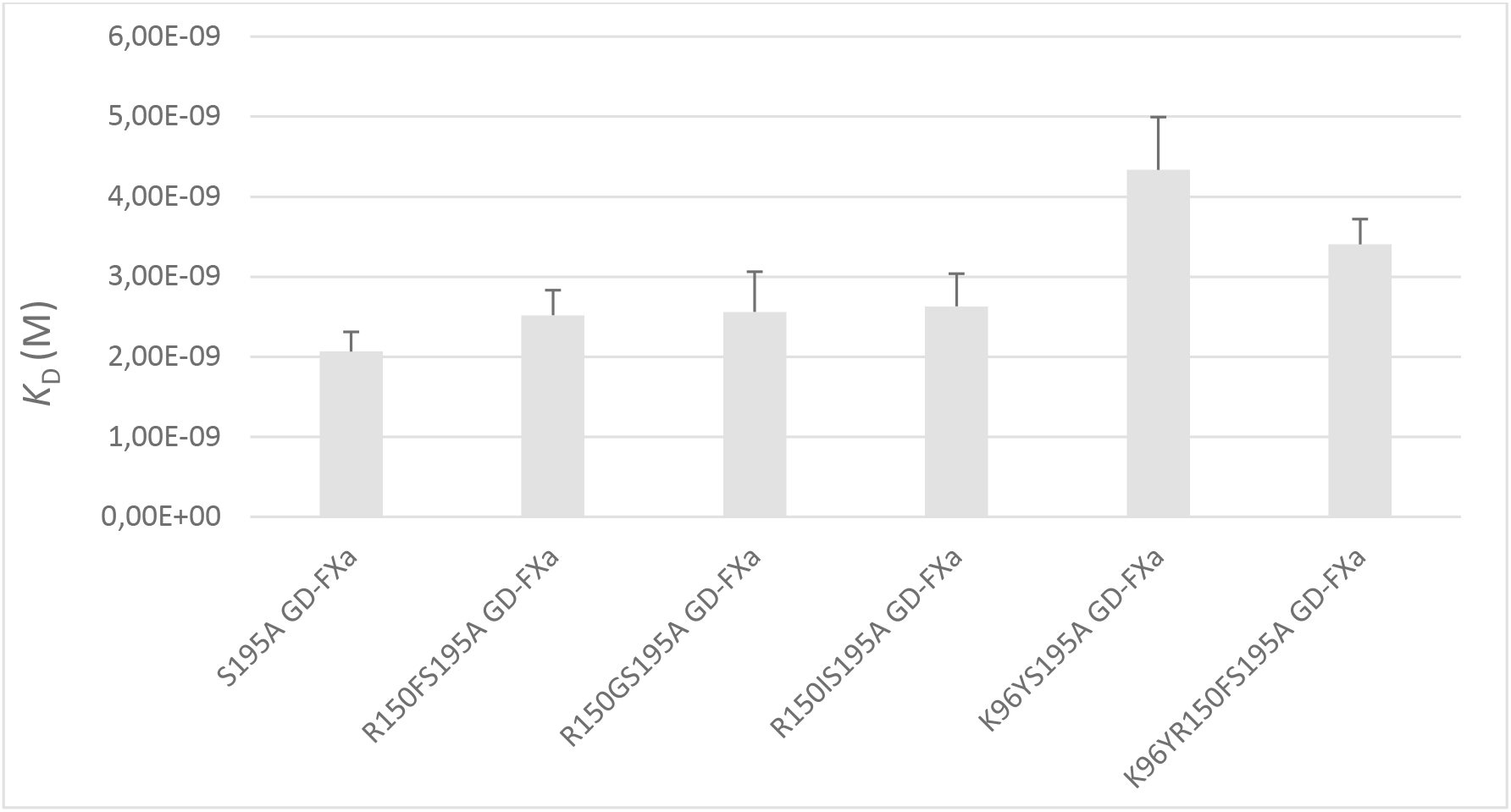
Comparison of the kinetic and dissociation constants of the S195A GD-FXa mutants with TFPI obtained from SPR experiments. Increasing concentrations of GD-FXa mutants (1.25–40 nM) were injected over immobilized TFPI (2600 and 4400 RU). The kinetic constants were calculated by fitting the data sets to a 1:1 binding model (see Supplementary Figure S2) and the ratio of the association (*k*_a_) and dissociation (*k*_d_) rate constants yielded the *K*_D_ constants. Values represent the means ± SD of two to four separate experiments.

The mutation R150F is the most efficient to improve TFPI binding and therefore the best candidate for improving S195A GD-FXa with respect to hemophilia treatment.

#### Thrombin generation assays

If GD-FXa mutants are more efficient TFPI traps than S195A GD-FXa, thrombin generation parameters should be consistent with computational data obtained by MD modeling. We performed thrombin generation assays in FVIII immuno-depleted plasmas spiked with different concentrations of mutants, all having the S195A mutation.

We compared two parameters from the thrombin generation assays: the thrombin peak height and the endogenous thrombin potential (ETP). As expected, the R150FS195A GD-FXa mutant having a higher association with TFPI is also more efficient in thrombin generation (Figure 3).

**Figure 3:**
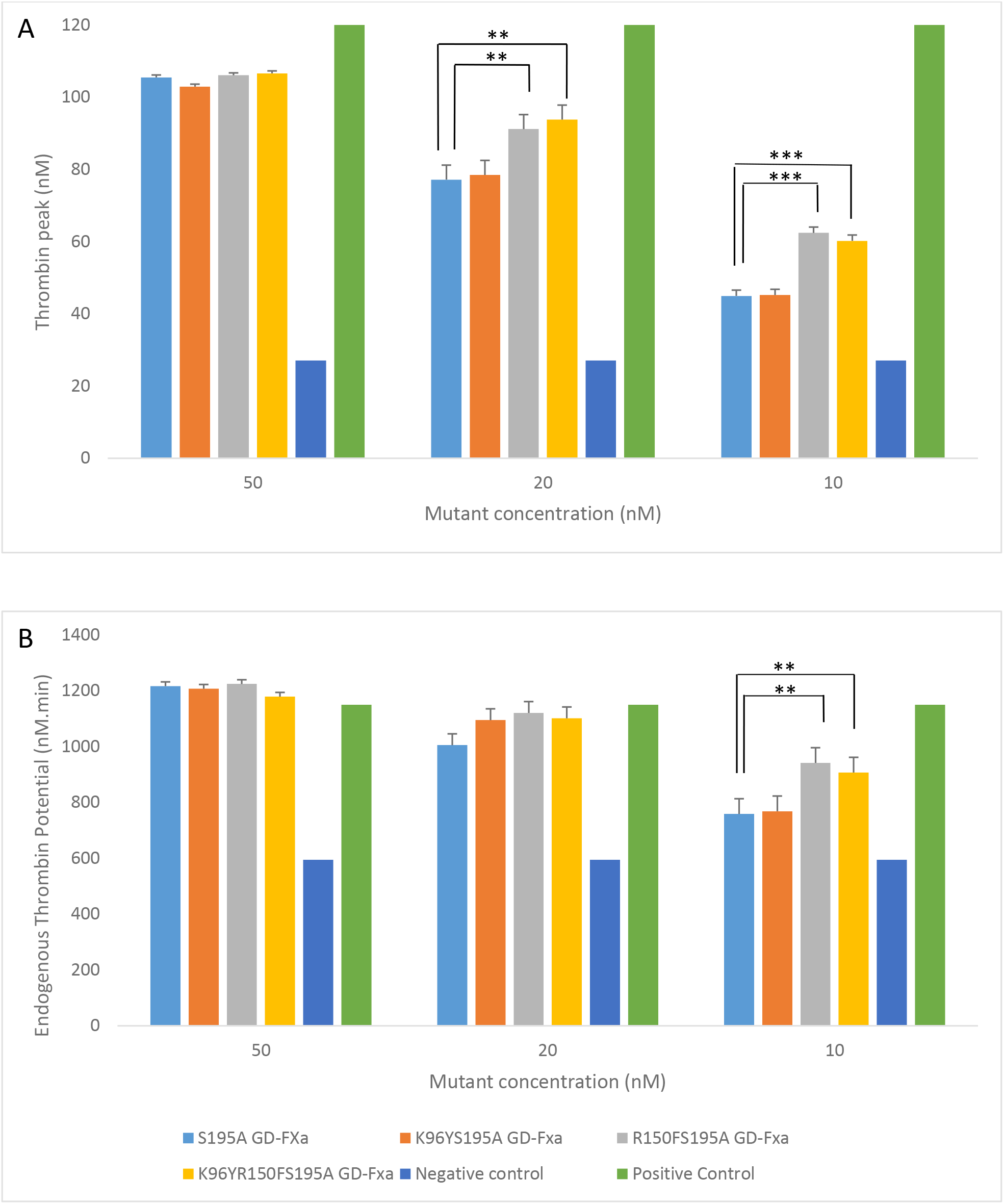
Thrombin generation assays. Plasma from severe hemophilia A patients was spiked with increasing concentrations of mutants in the presence of PPP-low reagent as described in Methods. The positive control was 1U/ml of recombinant Factor VIII in hemophilia A plasma (negative control). The curves were fitted using Thrombinoscope Software V5.0 (Diagnostica Stago) and (A) Thrombin peak height and (B) Endogenous thrombin potential (ETP) were calculated by the software. The bars represent the means +/- SD of 3 measurements. The statistical significance was calculated by two to two Tuckey confidence intervals (See Figure S5) and is depicted with stars (*: p<0.05; ** 0.02<p<0.01; ***: p<0.01)

For the ETP data, as can be seen in Figure 4, a marked favorable effect of R150FS195A mutant can be seen compared to S195A or K96YS195A mutant. This effect is seen only at low concentrations of 10 nM and is no longer significant at higher concentrations. On the contrary, the K96Y mutation had no effect, either alone in S195A GD-FXa or in combination with the R150F mutation (see Supplementary S4).

**Figure 4:**
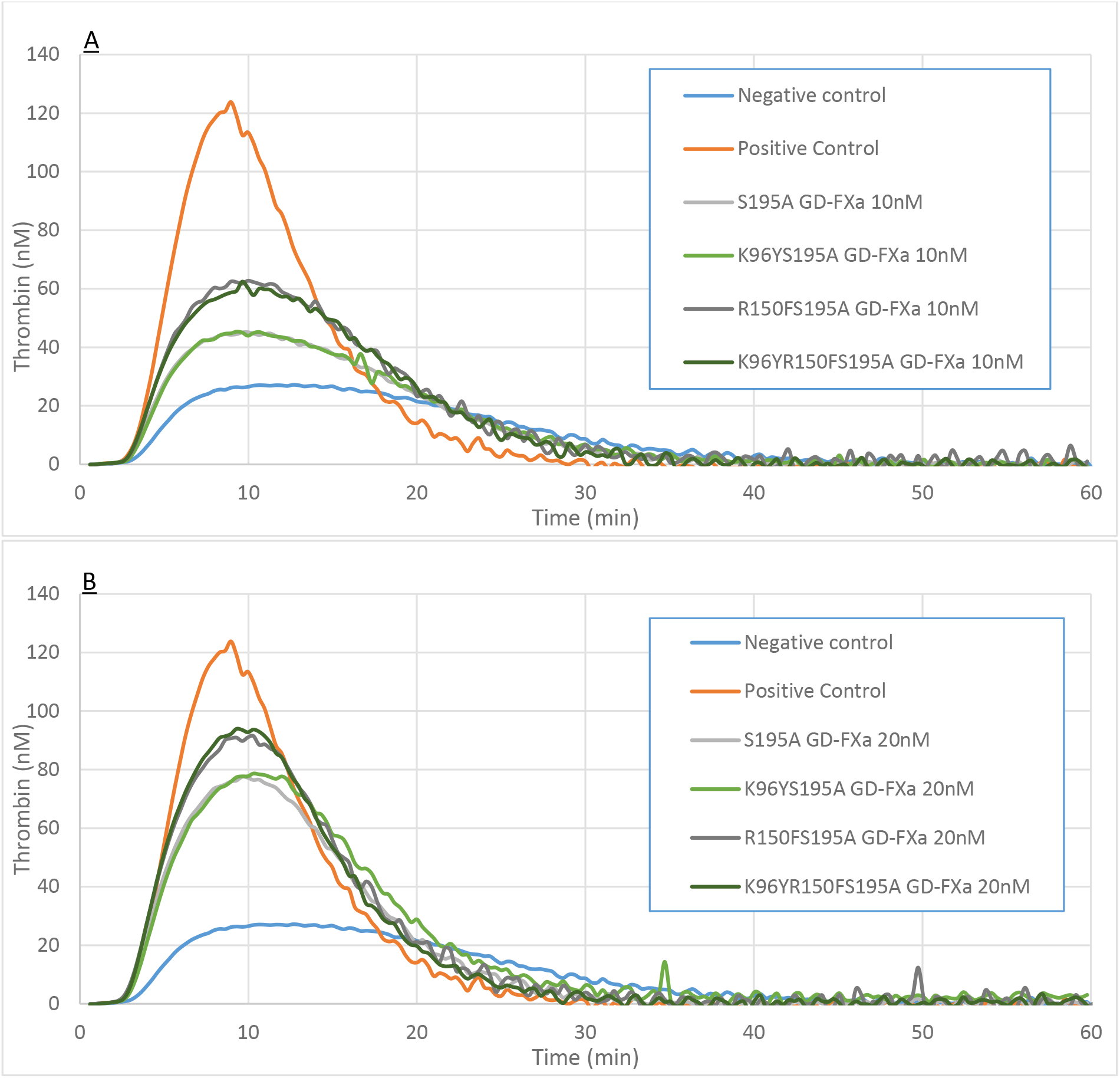
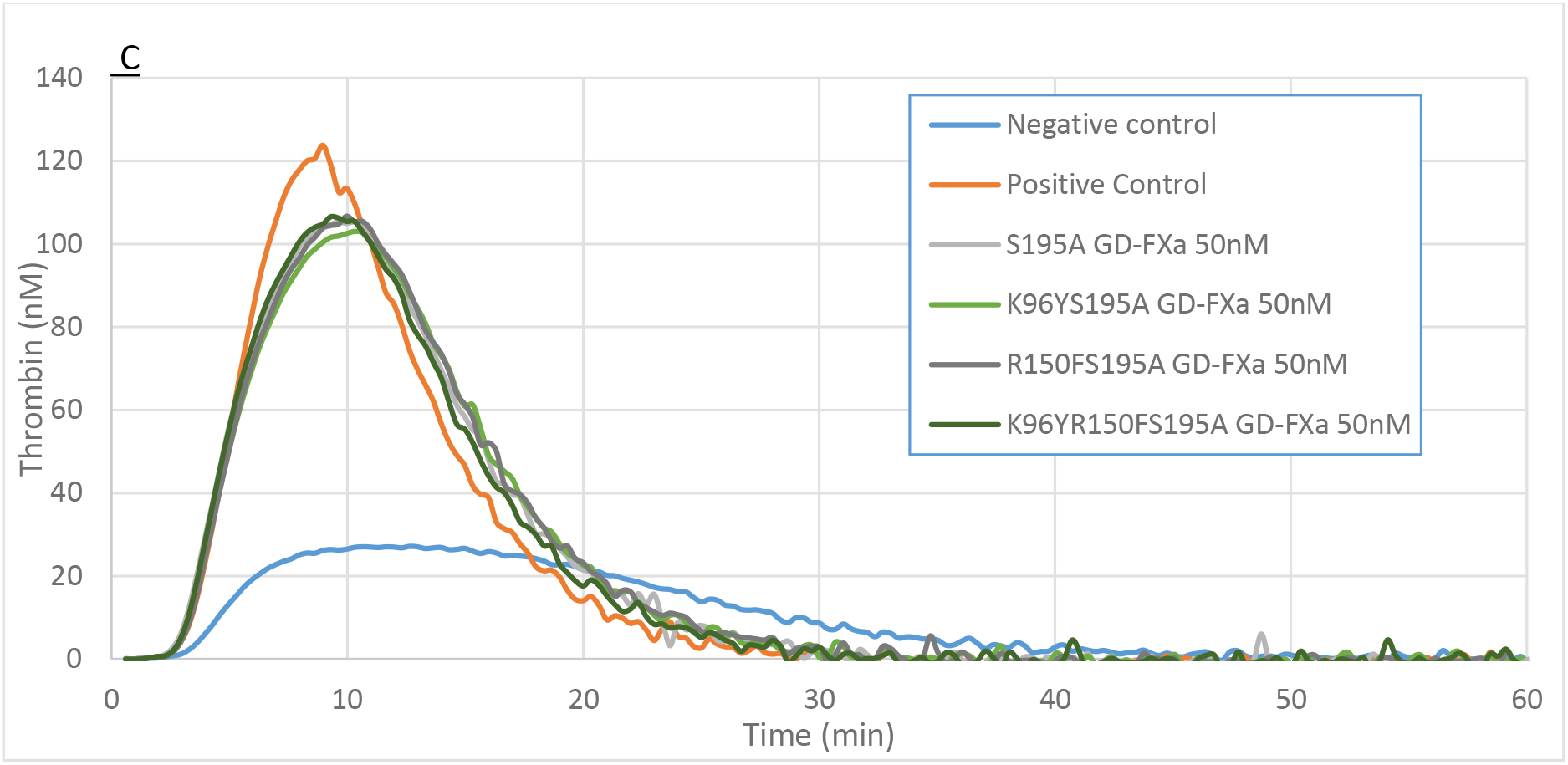
Graphs of Thrombin generation. The computed curves at the indicated concentrations of GD-FXa and mutants from the Thrombin generation assay triplicates (see Figure 3 for the calculated parameters) are plotted as a function of time, at different mutant concentrations. A: 10 nM; B: 20 nM; C: 50 nM.

The significant effect on ETP might reflect a longer half-life of the mutant that could be due to a reduced reactivity with AT. Conversely, at low concentrations, the effect of the R150F mutation on the thrombin peak height is significantly higher than that of S195A alone. In agreement with the binding data, the K96Y mutation did not have a favorable effect on the ETP nor on thrombin peak; furthermore, it counteracted the R150F mutation effect when both mutations were present. Thus, we observed a slight but significant favorable effect of the R150F mutation on both thrombin generation assay parameters, that paralleled the increased association with TFPI (lower *k*_a_).

#### SPR analysis of the interaction of GD-FXa mutants with antithrombin

Antithrombin belongs to the serpin family, it inhibits the serine protease activity of coagulation factors. Its mechanism of action involves covalent binding to the serine protease after cleavage of the reactive loop by the serine protease itself^23^. Catalytically inactive GD-FXa mutants cannot make covalent complexes, allowing their association and dissociation with immobilized AT to be analyzed by SPR. The binding curves were characterized by fast association, equilibrium and fast dissociation phases (Figure 5), allowing determination of the equilibrium dissociation constants using steady state analysis. The *K*_D_ value for the reference S195A GD-FXa mutant was 416 ± 42 nM (n=4); the K96YS195A GD-FXa mutant shows similar binding characteristics of 460 ± 76 nM (n = 4).

**Figure 5:**
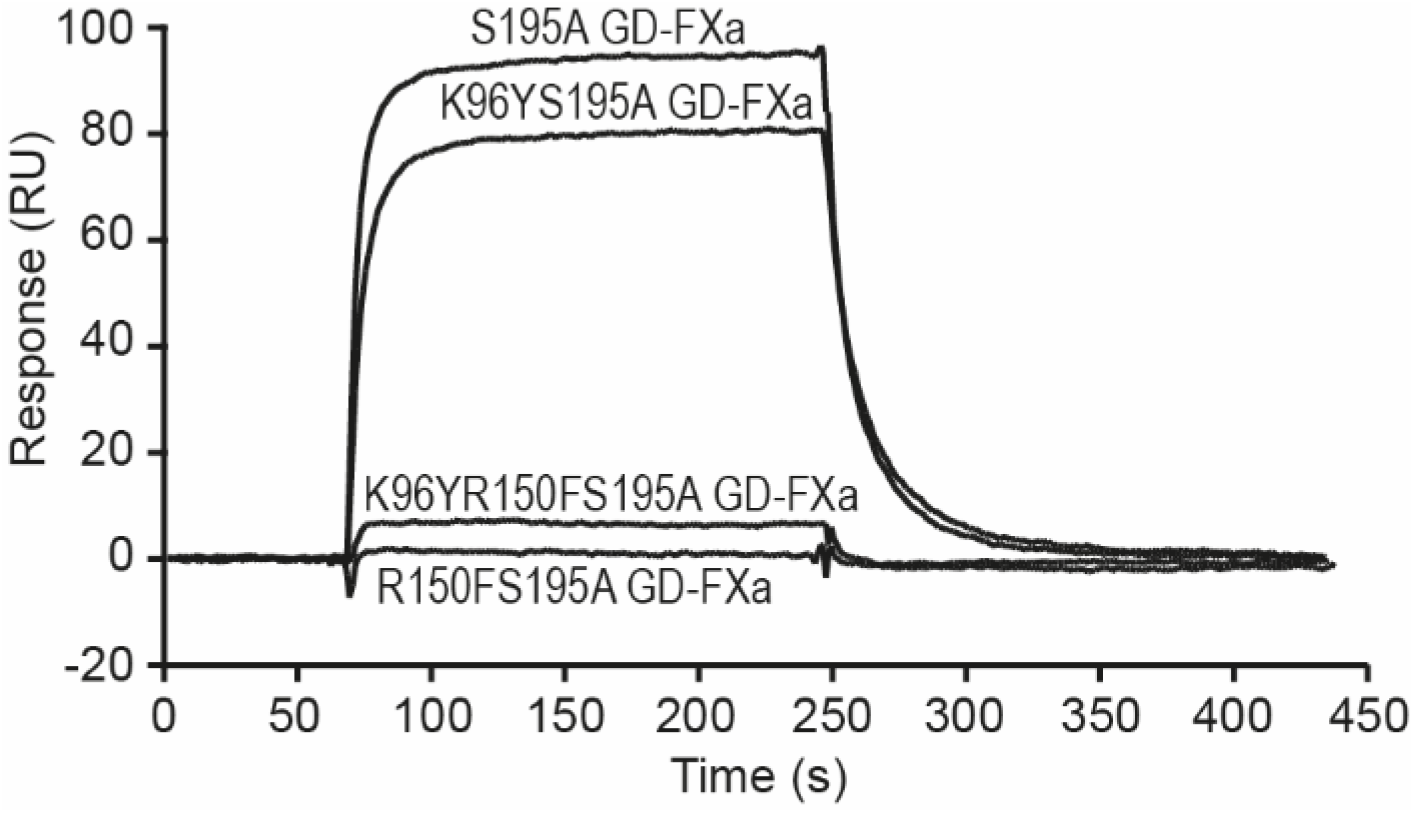
Representative analysis of the interaction of S195A GD-FXa mutants with immobilized antithrombin in the SPR experiments. The S195A GD-FXa mutants (500 nM) were injected over immobilized antithrombin (4400 RU) in 18 mM Hepes, 135 mM NaCl, 2.5 mM CaCl_2_, 0.005% surfactant P20, pH 7.35. Binding is expressed in resonance units (RU). Data are representative of 4 experiments on 3 different flow cells.

Not surprisingly from the structural analysis, in the case of the R150FS195A GD-FXa mutant, binding to AT was completely abolished, with only background signal observed that could not be analyzed by the software (Figure 5 and Supplementary Figure S3). The same was true for the other tested substitutions at the R150 position, namely R150G, R150I (see representative binding data sets for each mutant in Supplementary Figure S3).

Thus, the absence of AT binding for all engineered R150_FXa_ mutants was a good explanation for the increase in ETP observed in thrombin generation assays, in addition to better association with TFPI.

## Discussion

We have previously shown that GD-FXa and S195A GD-FXa proteins are procoagulant molecules targeting TFPI, that can be used as TFPI baits to restore coagulation in hemophilia patients^17^. S195A GD-FXa is also used as an antidote to DOACs to treat their accumulation that leads to the risk of bleeding^24–26^. In order to engineer GD-FXa variants with improved anti-TFPI activities, we performed theoretical molecular simulations to study the molecular interactions of GD-FXa with the TFPI-K2 domain. This allowed us to identify cold spots of interactions, i.e. residues that are unfavorable to the theoretical binding affinity between GD-FXa and TFPI-K2, and we rationally proposed mutations at those sites to enhance the TFPI-FXa interaction. Two cold spots of interactions were identified: K96_FXA_ and R150_FXa_ that were not involved in favorable local interactions. R150_FXa_ was mutated into a series of hydrophobic amino acids in order to improve local interactions with the facing tyrosine residue Y17_K2_. Among the engineered mutants, R150F_FXa_, introduced in the GD-FXa S195A background, improved the direct association with TFPI (*k*_a_) as monitored by SPR; concomitantly, higher levels of thrombin generation were achieved. Moreover, the mutation R150FFXa completely abolished the binding of the physiological inhibitor AT and this second biological activity is very likely to increase the variant bio-disponibility as well as the duration of thrombin generation (ETP).

The MD simulations predicted that the K96Y_FXa_ mutation might improve the thrombin generation parameters, which was not experimentally confirmed. This disagreement is probably due to the fact that we could only model the K2 domain of TFPI in the complex with GD-FXa, indeed, there is no experimental structure of the full-length TFPI nor of a larger TFPI fragment than a single Kunitz domain.

AT plays a critical role in regulating coagulation by forming a 1:1 covalent complex with thrombin and FXa^27, 28^. Thus, it is a target of choice for the development of therapeutic anti-hemophilic agents; as such, Fitusiran, a siRNA in clinical phase III, reduces AT production in people with hemophilia A or B, with and without inhibitors^29^. The AT interaction of the herein produced S195 GD-FXa variants was experimentally evaluated by SPR. The K96YS195A GD-FXA mutant shows a maintained interaction with AT as assessed by the equilibrium dissociation constants. This was expected as residue K96F_FXa_ is not in close proximity to AT in the ternary complex FXa-AT-fondaparinux, they are separated by more than 7Å in the crystal structure^23^. On the contrary, residue R150_FXa_ entertains strong hydrogen bonds with AT in the ternary complex; therefore, its substitution into a hydrophobic residue as envisaged from MD simulations to improve TFPI binding, was expected to reduce the stability of the AT-GD-FXa complex, and thus potentially prevent the derived FXa mutants from being trapped by AT.

The effect of the computationally identified R150F mutation on both the thrombin peak height and the ETP in Factor VIII immuno-depleted plasma is statistically significant. The engineered R150FS195A GD-FXa protein is a more efficient TFPI trap than S195A GD-FXa, illustrating that *in silico* predictions, combined with biological experiments can lead to designed proteins with improved properties. Work is in progress to design additional mutations with cumulative effects, on the R150FS195A GD-FXa scaffold, to lead to more powerful anti-TFPI agents and potentially nonreplacement hemophilic therapies.

## Acknowledgements

The project was funded by the Agence Nationale pour la Recherche 13-RPIB-0011 (AT, MCD, RM, LS, BP).

The simulations used the CIMENT infrastructure, which is supported by the Auvergne Rhône-Alpes region (CPER07_13 CIRA) and the Equip@Meso project (ANR-10-EQPX-29-01) of the program Investissements d’Avenir supervised by the Agence Nationale pour la Recherche. The authors would like to thank Pierre Girard for technical support. This work used the platforms of the Grenoble Instruct-ERIC Centre (ISBG; UMS 3518 CNRS-CEA-UJFEMBL) with support from the French Infrastructure for Integrated Structural Biology (FRISBI, ANR-10-INSB-05-02) and GRAL, a project of the University Grenoble Alpes graduate school (Ecoles Universitaires de Recherche) CBH-EUR-GS (ANR-17-EURE-0003) within the Grenoble Partnership for Structural Biology (PSB). We thank Isabelle Bally and Jean-Baptiste Reiser for assistance and access to the SPR facility. We are indebted to Xavier Brazzoloto who provided the drosophila expression system. We gratefully acknowledge our colleagues of LFB SA Toufik Abache, Alexandre Fontayne and Jean-Luc Plantier for fruitful discussions.

## Notes

The authors declare no conflict of interest

### Competing Interest Statement

The authors have declared no competing interest.

