## Supplemental Table S1 Figures S1 S2 S3 S4 for "Rationally designed Gla-domainless FXa as TFPI bait in hemophilia"

**Supplementary Data**

1. Molecular Dynamics of the complex between GD-FXa and the K2 domain of TFPI
2. Quality of the proteins used in the experiments
3. Kinetic analyses of the interaction of the S195A GD-FXa mutants with TFPI
4. Kinetic analyses of the interaction of the S195A GD-FXa mutants with antithrombin
5. Statistical analysis of Thrombin generation Assays
1. **Molecular Dynamics of the complex between GD-FXa and the K2 domain of TFPI**

The model of the complex between the catalytic chain of GD-FXa and the K2 domain of TFPI was derived from the crystal structures of FXa (PDB ID: 2J95^19^) and from the complex between porcine trypsin and human TFPI-K2 (PDB ID: 1TFX^20^).

The backbone atoms of the catalytic domains of both serine proteases were superimposed and the coordinates of FXa were then merged to the coordinates of the K2 domain of TFPI. As observed previously by other authors in a similar modeling study^20^ the resulting model of the complex showed collisions between the Y99_FXa_ side-chain and the disulfide bond Cys14-Cys38_K2_. In porcine trypsin, the smaller L99 does not face similar steric hindrance. This bump was relieved by orienting the Y99 side-chain towards the solvent, as performed by other authors^20^, thus favoring the formation of a hydrophobic cluster involving W215_FXa_, F174_FXa_ and I13_K2_, L39_K2_, C14_K2_ and C38_K2_.

The molecular dynamics (MD) simulations were performed with the CHARMM software^22^^,^^23^ using the CHARMM-36 parameters and topology sets including the CMAP correction^24^. The systems were fully solvated in a TIP3P^21^ water box of 16 Å edge larger than the protein dimensions. The molecular systems totalize about 35000 atoms including about 4580 protein atoms. Periodic boundary conditions were used to model the solvent. A minimization procedure was performed with decreasing harmonic forces on all the atoms, totalizing 3000 steps of steepest descent followed by 5000 steps of ABNR minimization without constraints. The structures were then subjected to a CPT molecular dynamics^34^ (1 atm, 298 K). The molecular systems were annealed from 48 K to 298 K over a period of 20 ps and maintained at that temperature with velocity scaling. The non-bonding interactions were considered up to a cutoff of 14 Å, and smoothly zeroed between 12 Å and 14 Å using a shifting potential. Covalent bonds involving hydrogens were kept fixed using the SHAKE algorithm (6), thus allowing an integration time step of 2 fs. The systems were equilibrated for 10 ns and the production runs lasted 100 ns. The trend of the overall potential energy, root mean square deviations (RMSD) and gyration radius, indicate that the systems are well stabilized after 10 ns. The analysis and visualization of the trajectory were performed using VMD^35^.

**Table S1**: Mean residual interaction energies of GD-FXa in the complex with the K2 domain (positive energies are highlighted in grey). A threshold of 0.1 kcal/mol in the energy was taken for clarity.

| **Residue Number** | **Mean Interaction**  **Energy (kcal/mol)** | **Standard Deviation**  **(kcal/mol)** |
| --- | --- | --- |
| 16 | **2,34** | **0,04** |
| 17 | **0,32** | **0,01** |
| 18 | **-0,07** | **0,00** |
| 32 | **0,16** | **0,00** |
| 33 | **-0,66** | **0,01** |
| 34 | **0,84** | **0,02** |
| 35 | **5,79** | **0,19** |
| 36 | **-0,91** | **0,30** |
| 37 | **-64,47** | **1,58** |
| 38 | **0,31** | **0,03** |
| 39 | **-78,34** | **2,18** |
| 40 | **-5,69** | **0,13** |
| 41 | **-7,95** | **0,20** |
| 42 | **-3,34** | **0,06** |
| 43 | **-0,27** | **0,01** |
| 55 | **-0,02** | **0,01** |
| 56 | **-0,22** | **0,01** |
| 57 | **-5,30** | **0,18** |
| 58 | **-2,79** | **0,08** |
| 59 | **-0,40** | **0,03** |
| 60 | **-2,67** | **0,19** |
| 61 | **-18,25** | **0,43** |
| 61A | **-1,42** | **0,07** |
| 62 | **-33,28** | **1,81** |
| 63 | **-1,80** | **0,23** |
| 90 | **0,11** | **0,01** |
| 94 | **-0,23** | **0,04** |
| 96 | **0,37** | **0,12** |
| 138 | **0,44** | **0,01** |
| 140 | **-0,10** | **0,00** |
| 141 | **-0,31** | **0,01** |
| 142 | **-0,75** | **0,02** |
| 143 | **-9,83** | **0,47** |
| 145 | **0,39** | **0,01** |
| 147 | **-2,09** | **0,13** |
| 148 | **-13,22** | **1,31** |
| 150 | **0,95** | **0,15** |
| 151 | **-5,28** | **0,47** |
| 152 | **-0,36** | **0,02** |
| 158 | **0,15** | **0,00** |
| 159 | **-0,02** | **0,00** |
| 160 | **0,30** | **0,00** |
| 172 | **0,10** | **0,00** |
| 174 | **-1,03** | **0,04** |
| 181 | **-0,14** | **0,00** |
| 182 | **0,39** | **0,01** |
| 183 | **-1,29** | **0,02** |
| 184 | **0,46** | **0,01** |
| 186 | **-0,21** | **0,00** |
| 187 | **0,33** | **0,02** |
| 188 | **-0,22** | **0,01** |
| 189 | **-60,61** | **0,36** |
| 190 | **-5,05** | **0,10** |
| 191 | **-2,66** | **0,06** |
| 192 | **-18,66** | **0,24** |
| 193 | **-7,08** | **0,05** |
| 194 | **-4,40** | **0,07** |
| 195 | **-3,99** | **0,08** |
| 196 | **0,13** | **0,00** |
| 198 | **0,36** | **0,01** |
| 199 | **0,65** | **0,02** |
| 213 | **-0,19** | **0,05** |
| 214 | **-2,44** | **0,07** |
| 215 | **-7,72** | **0,13** |
| 216 | **-7,84** | **0,23** |
| 217 | **-7,56** | **0,16** |
| 219 | **-16,38** | **0,11** |
| 220 | **1,60** | **0,04** |
| 221 | **3,24** | **0,04** |
| 222 | **-0,77** | **0,41** |
| 223 | **-0,14** | **0,00** |
| 223A | **0,11** | **0,00** |
| 224 | **0,58** | **0,04** |
| 225 | **0,80** | **0,03** |
| 226 | **2,76** | **0,04** |
| 227 | **-1,71** | **0,03** |
| 228 | **-1,22** | **0,07** |

2. **Quality of the proteins used in the experiments**

**A**
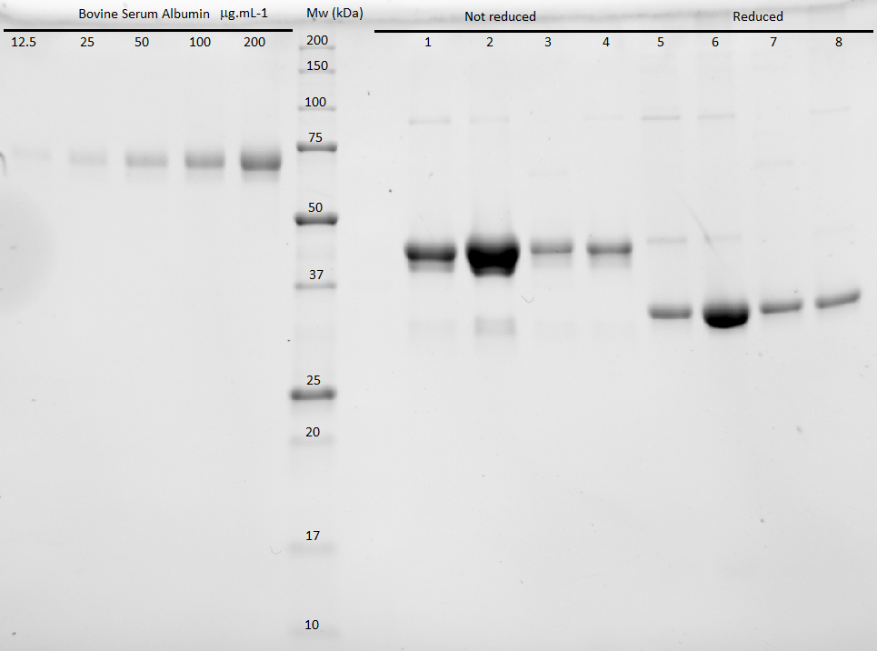

**B**
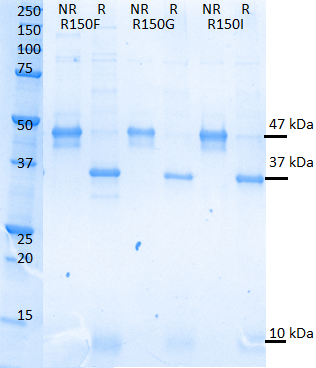

**Figure S1: Purity of recombinant proteins used in the study**

Samples of purified GD-FXa mutants, all containing the S195A mutation, were either reduced (R) or not reduced (NR) with 50 mM dithiothreitol to separate the light chain and the heavy chain and loaded on the Stain Free gel (Bio-rad). Since the light chain does not contain tryptophan, it is not visualized by the Bio-rad Stain Free procedure. (A) Mutants used in Thrombin generation assays: K96YS195A (lanes 1,5), S195A (lanes 2, 6), R150FS195A (lanes 3, 7), K96YR150FS195A (lanes 4, 8). The calibration quantities of bovine serum albumin are shown on the left. (B) For the other mutants studied by SPR, gels were stained with Coomassie Blue. The molecular weight of the chains is indicated.

3. **Kinetic analyses of the interaction of the S195A GD-FXa mutants with TFPI.**

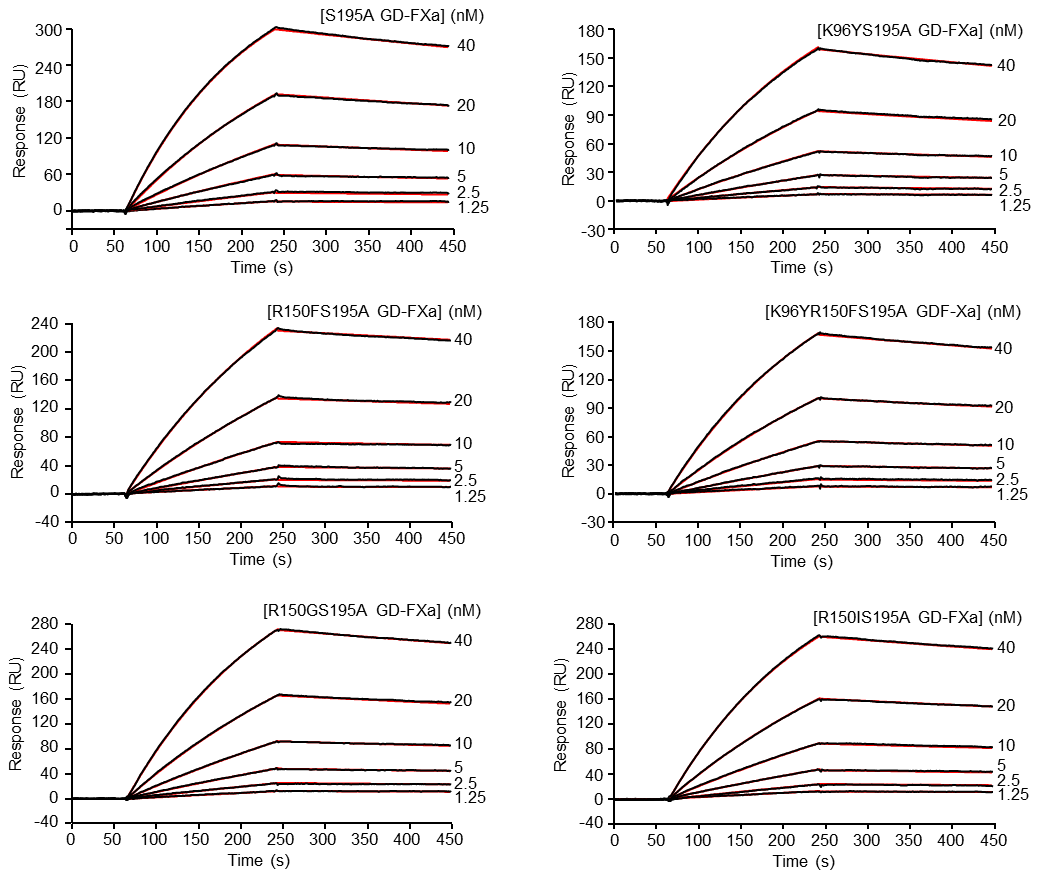

**Figure S2. Kinetic analyses of the interaction of the S195A GD-FXa mutants with immobilized TFPI from SPR experiments**Sixty µl of the S195A GD-FXa mutants at the indicated concentrations were injected over immobilized TFPI (1500 RU) in 18 mM Hepes, 135 mM NaCl, 2.5 mM CaCl_2_, 0.005% surfactant P20, pH 7.35 at a flow rate of 20 µl/min. The binding signals shown were obtained by subtracting the signal over the reference surface and further subtraction of buffer blanks. Fits are shown as red lines and were obtained by global fitting of the data using a 1:1 Langmuir binding model. Each kinetic analysis shown is representative of two or three experiments on separate flow cells.

4. **Kinetic analyses of the interaction of the S195A GD-FXa mutants with AT.**

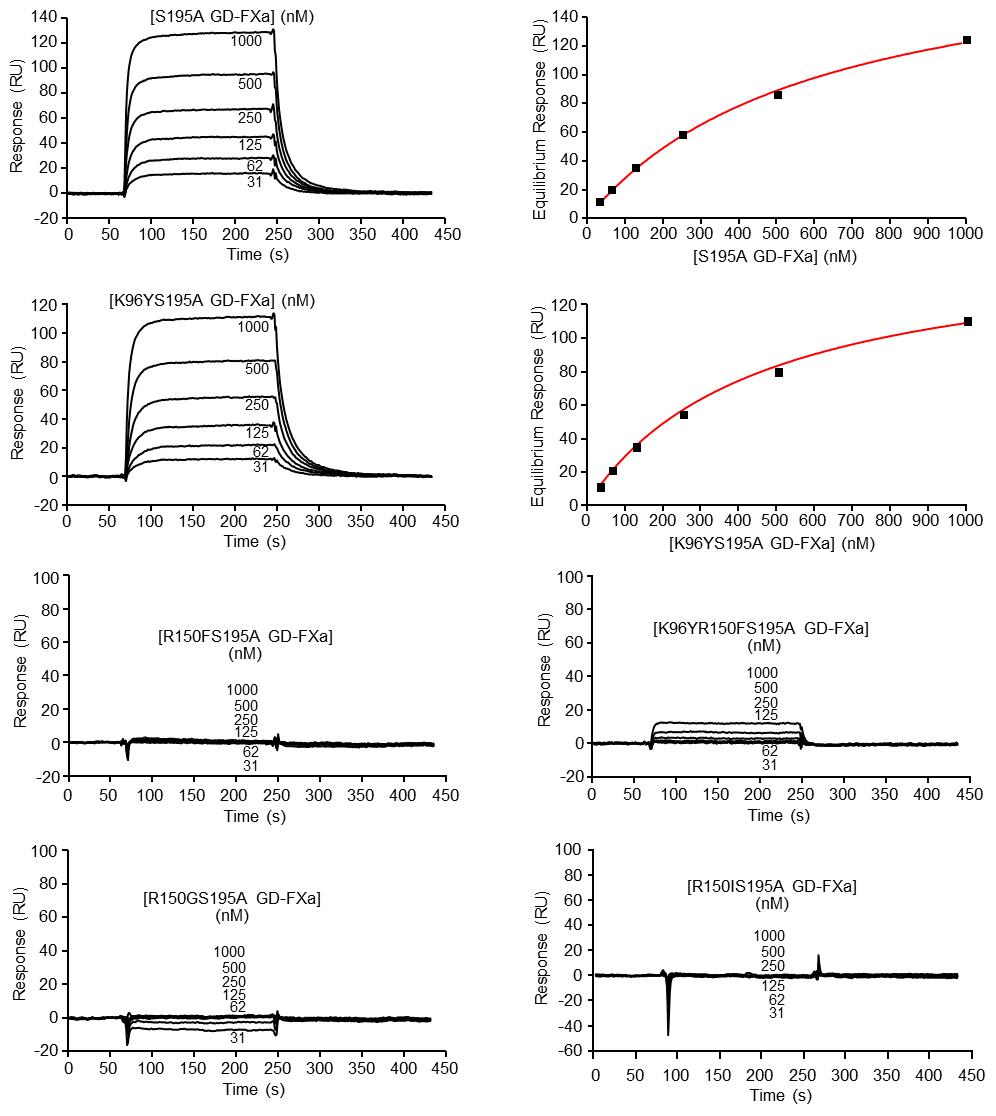

**Figure S3. Kinetic analyses of the interaction of the S195A GD-FXa mutants with immobilized AT from SPR experiments.** Sixty µl of the S195A GD-FXa mutants at the indicated concentrations were injected over immobilized AT (4400 RU) in 18 mM Hepes, 135 mM NaCl, 2.5 mM CaCl_2_, 0.005% surfactant P20, pH 7.35 at a flow rate of 20 µl/min. The binding signals shown were obtained by subtracting the signal over the reference surface and further subtraction of buffer blanks. Fits shown as red lines were obtained using steady state analysis. Each kinetic analysis or set of curves shown is representative of two to four experiments on separate flow cells

5. **Statistical analysis of Thrombin generation Assay**

### **Mutant pairwise comparisons**

For each response, a one-way ANOVA procedure was carried out for each concentration (10, 20 and 50 nM) to quantify the difference between the mutant types. When a significant difference resulting from the ANOVA was found (p<0.05 level), a Tukey HSD post-hoc test was performed to determine which specific mutant group means (compared to each other) were different. The test compared all possible pairs of averages.

For the ETP response (top), no statistically significant difference was found between averaged responses as determined by one-way ANOVA at 20 and 50 nM concentration (p-value = 0.163 and p-value = 0.815 respectively). However, a statistically significant difference was observed at 10 nM (p-value = 0.005). Therefore, Tukey confidence interval tests were performed and showed statistically significant differences between the mutants containing R150F or not were observed at 10 nM (Figure S4a).

For the thrombin peak height response (bottom), a statistically significant difference was observed by one-way ANOVA at 10 and 20 nM (with respective p-values equal to 0.001 and 0.004) but not at 50 nM (p-value = 0.591). Simultaneous Tukey confidence interval tests were computed for the thrombin peak height at 10 nM (Figure S4b) and 20 nM (Figure S4c).

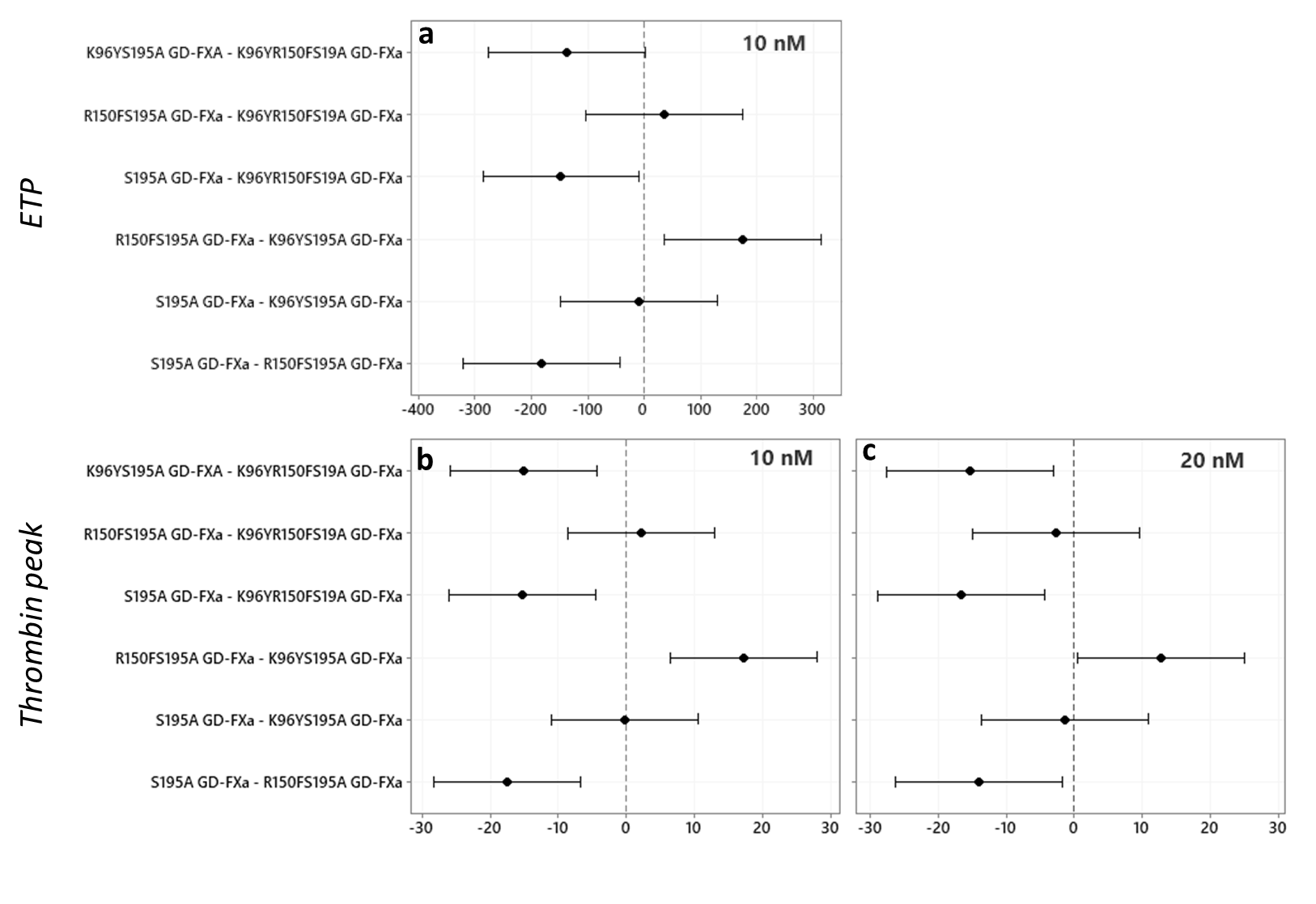

**Figure S4: Post-Hoc Tukey simultaneous 95% confidence interval.**

The confidence intervals are drawn for each mutant pair. If an interval does not contain the zero, the corresponding average values are significantly different at the p<0.05 level.
